# Mitigating assembly and switch errors in phased genomes of polar fishes reveals haplotype diversity in copy number of antifreeze protein genes

**DOI:** 10.1101/2025.09.09.675170

**Authors:** Owen W. Moosman, Joanna L. Kelley, Samuel N. Bogan

**Affiliations:** University of California, Santa Cruz, Department of Ecology and Evolutionary Biology; University of California, Santa Cruz, Department of Molecular, Cell, & Developmental Biology

**Keywords:** Genome assembly, copy number, pangenomes, antifreeze proteins, adaptation, structural variation

## Abstract

Phased genomes and pangenomes are enhancing our understanding of genetic variation. Accurate phasing and assembly in repetitive regions of the genome remain challenging, however. Addressing this obstacle is crucial for studying structural genomic variation such as copy number variations (CNVs) common to repetitive regions. Polar fishes, for example, evolved repetitive tandem arrays of antifreeze protein (AFP) genes that facilitated adaptation to freezing and expanded in copy number in colder environments. AFP CNVs remain poorly characterized in polar fishes and may be illuminated by haplotype-aware approaches. We performed long-read sequencing of two polar fishes in the suborder Zoarcoidei and leveraged published long-read data to assemble phased genomes. We developed a workflow to measure haplotype diversity in CNV while controlling for misassembly and switch errors—a change from one parental haplotype to another in a contiguous assembly. We present *gfa_parser*, which computes and extracts all possible contiguous sequences for phased or primary assemblies from graphical fragment assembly files, and *switch_error_screen*, which flags potential switch errors. *gfa_parser* revealed that assembly uncertainty was ubiquitous across AFP array haplotypes and that standard processing of graphical fragment assemblies can bias measurement of haplotype CNVs. We detected no switch errors in AFP arrays. After controlling for misassembly and switch error, we detected haplotype diversity of AFP CNVs in all studied polar Zoarcoidei species and in 60% of AFP arrays. Intraindividual haplotype diversity spanned differences of 3–16 copies. Our workflow revealed intraspecific genomic diversity in zoarcoids that likely fueled evolution of AFP copy number across temperature.

## 1. Introduction

Advances in long-read sequencing and genome assembly have greatly improved our ability to detect and analyze genetic diversity. These advances include phased genomes, which distinguish between paternal and maternal inherited haplotypes (full phasing) or distinct haploid sequences (pseudo-phasing), and pangenomes, which contain the full diversity of haplotypes across samples of a species or population (Browning and Browning, 2011; Eizenga *et al*., 2020; Porubsky *et al*., 2021). Pangenomic approaches have enabled detecting and phasing of large structural genomic variants (SVs) that are longer than what can be detected using alignment of resequenced long-read data (Ebert *et al*., 2021). These SVs include large translocations, inversions, insertions, deletions, and copy number variations (CNVs). To date, most haplotype-aware phased genomes or pangenomes have been assembled for model species with trio sequence data from the maternal, paternal, and offspring genomes (Koren *et al*., 2018; Chaisson *et al*., 2019). However, some of the most valuable systems for investigating haplotype diversity, particularly among SVs, are natural populations. Many natural populations must be sampled in the field where collecting trio samples is difficult, if not impossible. Phasing can be performed in these cases using chromatin interaction data such as Hi-C sequence data (Kronenberg *et al*., 2021) or physical phasing inferred from overlapping reads (Cheng *et al*., 2021), but chromatin interaction phasing can be less accurate compared to trio sequencing (Kronenberg *et al*., 2021) and also presents feasibility issues during *in situ* sampling (Whibley *et al*., 2021).

In addition to understanding SVs, successful phasing can also improve assembly accuracy in repetitive regions that are difficult to resolve (Xu *et al*., 2021; Porubsky *et al*., 2023), and where SVs such as CNVs are more common (Carvalho and Lupski, 2016; Collins and Talkowski, 2025). However, trio phasing, chromatin-interaction phasing, and physical phasing are each difficult in repetitive regions (Kronenberg *et al*., 2021; Hu *et al*., 2024). When using haplotype-phased data from natural populations to study SVs, artifacts of genome misassembly and switch errors, a change from one parental allele to another in a contiguous segment of the genome assembly (contig), may appear to be SVs (Fig. 1). Artifacts and switch errors are common in repetitive regions (Choi *et al*., 2018). Unfortunately, there is a paucity of tools that allow us to distinguish genome misassembly and phasing errors from actual variation between haplotypes in phased genomes and pangenomic data.

**Figure 1.**
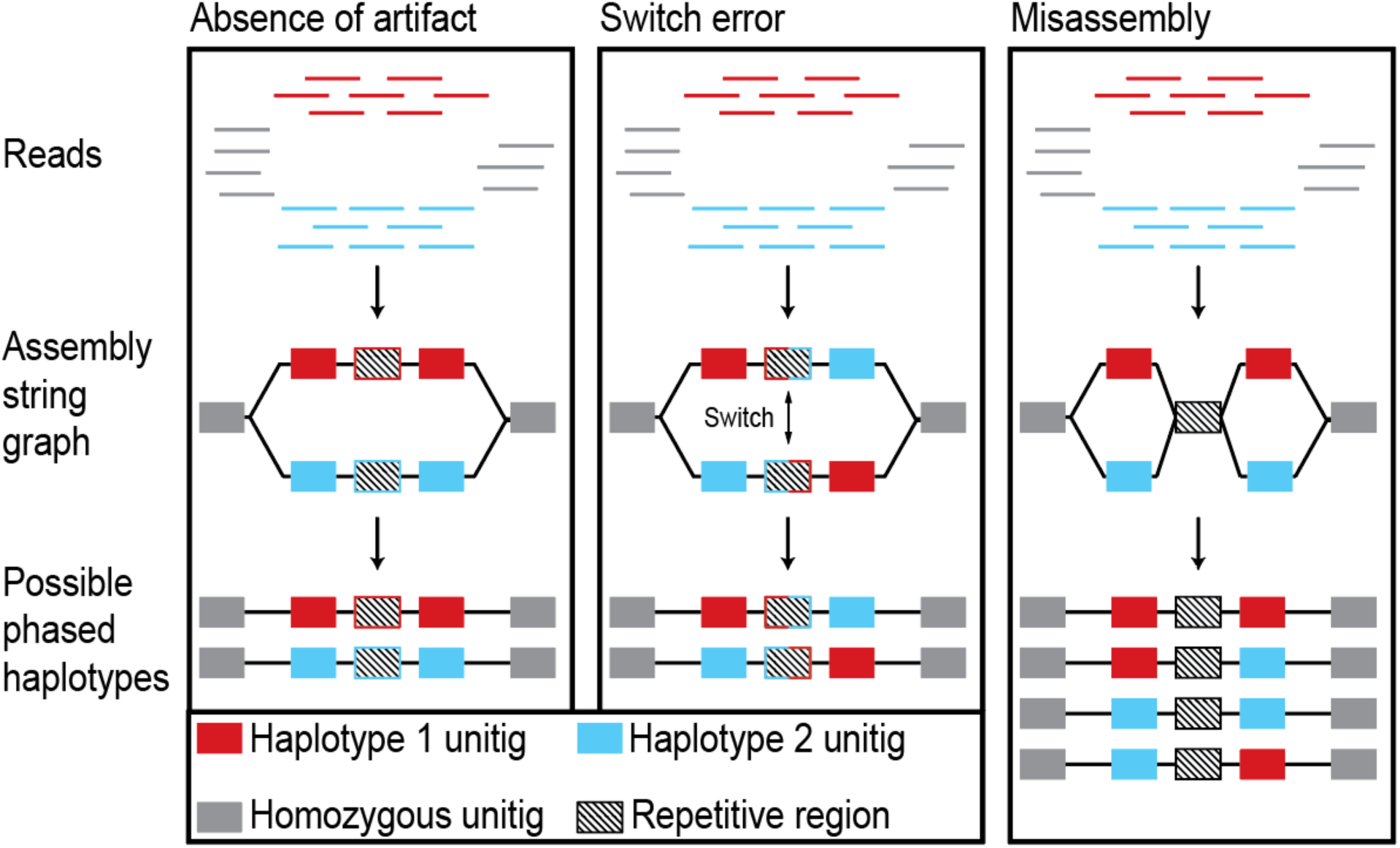
Diagram of genome assemblies with an absence of artifact and two possible assembly artifacts: switch error and misassembly. Each box represents a set of possible outcomes during phased assembly. Raw reads colored by unique haplotype of origin are organized into assembly string graphs. Haplotypes are represented by colored blocks in heterozygous regions creating assembly “bubbles”. Colored borders represent haplotype of origin in repetitive regions. Black-bordered unitigs represent those that are present in both haplotype assemblies. String graphs are phased into distinct haplotype-specific assemblies.

Recently, phased genomes have revealed CNVs between haplotypes tied to silk production in insects and spiders (Frandsen *et al*., 2023), immune function in pearl oysters (Takeuchi *et al*., 2022), and stress response in paperbark trees (Chen *et al*., 2022). These studies yield critical insights, but were unable to evaluate the influence of assembly or switch errors on the inference of haplotype diversity. This is largely because approaches for doing so have not been reported. Information about assembly uncertainty (a metric for the likelihood of misassembly) and heterozygosity between haplotypes of a genome are contained within graphic fragment assembly (GFA) files produced by popular assembly packages such as hifiasm (Cheng *et al*., 2021), Shasta (Shafin *et al*., 2020), and Canu (Koren *et al*., 2017). Bioinformatic tools that use GFA files to compute and extract all possible assemblies for a given haplotype would enable the validation of haplotype diversity in SVs against misassembly artifacts in repetitive or complex genomic regions (Fig. 1).

The causes and consequences of switch errors during genome assembly are well documented (Majidian and Sedlazeck, 2020; Holt *et al*., 2024; Nie *et al*., 2024), but their downstream effect on measurements of CNVs has received little attention. Switch errors can arise from misassembly of heterozygous regions (Fig. 1) or incorrect phasing of parental alleles during assembly. While many genome assembly algorithms aim to reduce the risk of switch errors (Martin *et al*., 2016; Rhie *et al*., 2020; Cheng *et al*., 2021), computational tools for detecting switch errors in phased genomes are limited.

The genome assembler hifiasm is a popular tool for assembling high accuracy Pacific Biosciences (PacBio) HiFi reads. A summary of the assembly and phasing processes of hifiasm is shown in Figure 1. First, raw HiFi reads are assembled into a GFA composed of a network of unitigs—a unit of assembly representing a block of reads with no alternative overlaps. GFA networks are composed of unitig nodes and edges that represent partial overlaps between unitigs (Myers, 2005). Contigs, by contrast, are generated from strings of partially overlapping unitigs. In a GFA, heterozygous regions form ‘bubbles’ which diverge from a common homozygous unitig sequence into two different unitigs representing distinct haplotypes. Raw GFA files undergo purging of some unresolved regions and ‘bubble popping’, a process of removing alternative unitigs at small bubbles attributed to somatic mutation or sequencing error. The new, purged graph is then phased into two distinct haplotype assemblies according to remaining haplotype bubbles (Cheng *et al*., 2021). In Figure 1, a normal assembly is shown alongside theoretical assemblies with switch errors and misassembly, which create alternative phased haplotype assemblies with incorrect resolution of the reads.

One system with well-documented gene contraction and expansion is type III antifreeze proteins (AFPs) in the suborder of polar fish Zoarcoidei. Zoarcoidei evolved AFPs that have allowed them to adapt to polar environments across the globe, including both the Arctic and Southern Oceans (Hobbs *et al*., 2020; Hotaling *et al*., 2021; Bogan *et al*., 2024). The AFP III protein functions by binding to nucleating ice crystals in serum and attenuating their expansion. The result of this activity is a lowering of cellular freezing point through thermal hysteresis, an increase in the thermal distance between freezing and melting point (Jia *et al*., 1996). AFP III evolved from neofunctionalization of the enzyme sialic acid synthase B, maintaining the first and sixth exons which now make up the signaling peptide and ice-binding domain of the AFP III (Deng *et al*., 2010). In some species, translocation-duplication resulted in AFP arrays on multiple chromosomes that are distinct from the ancestral AFP array (Bogan *et al*., 2024). The ancestral array is identifiable due to its conservation of synteny among multiple genes that flank the array as shown in Figure 2 (Deng *et al*., 2010; Hotaling *et al*., 2023). Increased copy number of AFPs in Zoarcoidei may have facilitated adaptation to colder temperatures in shallow-water species (Desjardins *et al*., 2012; Bogan *et al*., 2024). However, the amount of intraspecific variation or haplotype diversity in AFP copy number has not been tested at a large scale in any polar fish.

**Figure 2.**
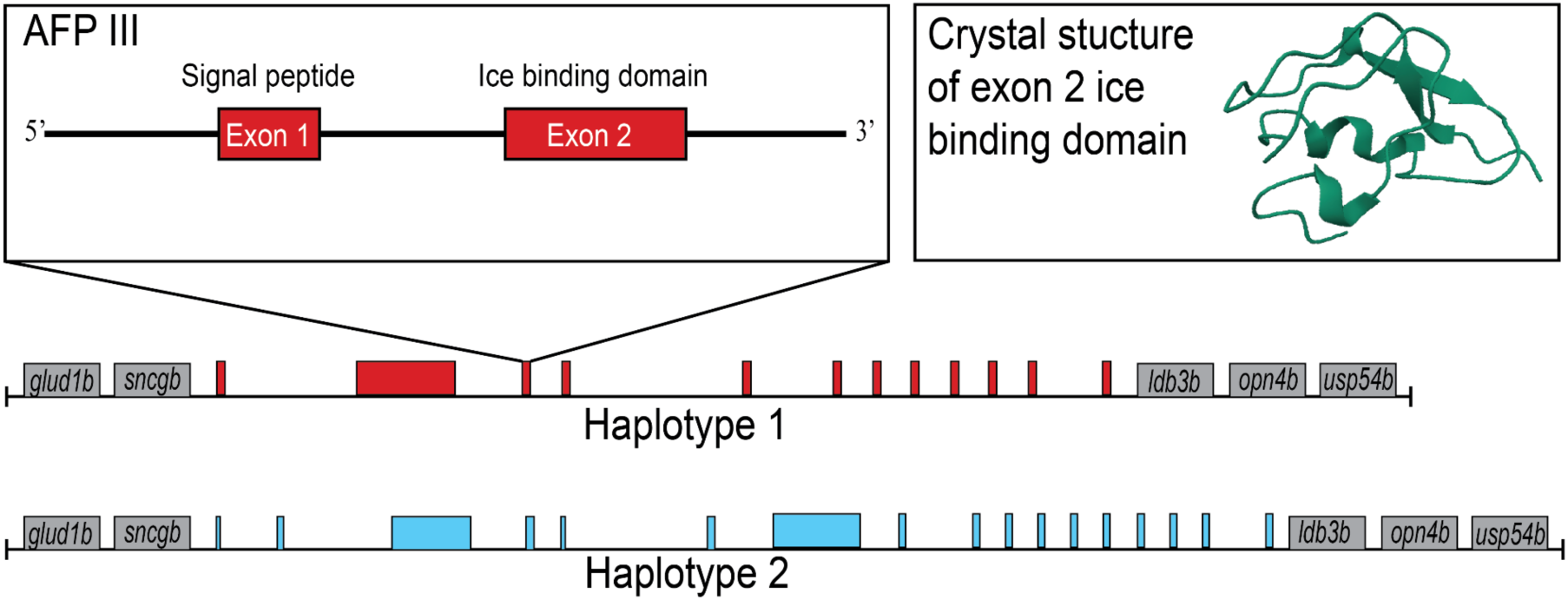
Type III antifreeze protein and tandem array structure. Theoretical representation of tandem repeat AFP array along homologous chromosomes is shown alongside the exon structure of AFP III gene and crystal structure of AFP III protein (Ye *et al*., 1998). AFP copies are represented by colored boxes and are colored by haplotype. Upstream and downstream flanking genes are represented in gray, which are syntenic at the ancestral AFP III array across Zoarcoidei.

Our goals for this study were two-fold. First, to develop and test new computational tools for evaluating the effects of misassembly and switch errors on analyses of structural variation in phased assemblies. We confirmed the applicability of these tools to assemblies and GFAs produced by hifiasm (Cheng *et al*., 2021), Shasta (Shafin *et al*., 2020), Verkko (Rautiainen *et al*., 2023), and minigraph (Li *et al*., 2020). Second, to apply these tools to determine the extent of haplotype diversity and heterozygosity in AFP copy number across Zoarcoidei, a system for studying environmental adaptation that currently lacks information on genomic diversity. We devised two methods to quantify the effects of assembly and phasing artifacts and distinguish artifacts from actual variation in AFP copy number between haplotypes. The reliability of assembly and phasing of the AFP III array was investigated by quantifying the variance in copy number estimates due to misassembly using assembly string graphs and screening for switch errors among haplotypes. By performing PacBio HiFi sequencing on two species of polar zoarcoids and compiling published HiFi data across Zoarcoidei, we applied this approach to phased genome assemblies to measure haplotype diversity in AFP CNVs.

## 2. Methods and Materials

### 2.1. Sampling, long-read genome sequencing, and additional data acquisition

Wild-caught individuals of the Antarctic and temperate-Arctic zoarcoids *Lycodichthys dearborni* and *Zoarces americanus* were collected and sequenced for phased *de novo* genome assembly. *L. dearborni* were collected via baited fish trap in Cape Evans of the McMurdo Sounds, Ross Sea (−77.63 °S, 166.41°E) in November 2011. High molecular weight DNA was extracted from the spleen and heart of a single individual using the Mag-Bind® Blood & Tissue DNA HDQ 96 Kit (Omega-BioTek, Norcross, GA, USA). DNA extractions underwent additional purification and concentration using the Zymo Research (Irvine, CA, USA) Genomic DNA Clean & Concentrator®-10 kit, which was followed by short-read elimination using the Circulomics (Baltimore, MD, USA) Short Read Eliminator Kit. Library preparation was performed separately for both extractions using the PacBio SMRT-Bell v3 kit without DNA shearing. Each library was separately sequenced on PacBio Revio flow cells (PacBio, San Jose, CA, USA) using a 30-hour movie time.

*Zoarces americanus* was collected via bottom trawl by the Virginia Institute of Marine Science NorthEast Area Monitoring and Assessment Program (Bonzek, 2012; Saba *et al*., 2023) off the coast of Newport, RI, USA, offshore of the mouths of Narragansett and Buzzards Bay, Cape Cod, USA (41.37 °N, −71.11°W) in May 2024. High molecular weight DNA was extracted from spleen dissected from a single individual using the PacBio Nanobind PanDNA Kit tissue extraction protocol followed by treatment with the Circulomics Short Read Eliminator Kit. Extracted DNA was sheared using tagmentation, which cleaved DNA to a modal length of 19.9 kilobases (kb) via insertion of internal adapter sequences by tagmentase (see Supplemental Methods). Library preparation was performed using the PacBio SMRT-Bell v3 kit. The library was size-selected for >7 kb via LightBench (Yourgene Health; Manchester, UK). The library was sequenced on a PacBio Revio flow cell using a 30-hour movie time. HiFi binary alignment map (BAM) files for *L. dearborni* and *Z. americanus* were converted to FASTQ using bedtools bam2fastq (Quinlan and Hall, 2010). Read and contig N50 were calculated for both species using SeqKit stat v2.5.1 (Shen *et al*., 2024).

FASTQ files containing HiFi reads were downloaded from the NCBI Sequence Read Archive (SRA) for *Leptoclinus maculatus* (SRR25603844; SRR25603845), and *Pholis gunnellus* (ERR6436364; ERR6412365) (Programme *et al*., 2022); using prefetch and fasterq-dump from sratoolkit v3.0.0 (Leinonen *et al*., 2011). Raw HiFi reads for the wolffishes *Anarhichas lupus* (PRJNA980960) and *Anarhichas minor* (PRJNA982125) were obtained from Bogan et al. (Bogan *et al*., 2024).

### 2.2. Genome assembly and annotation

Haplotype-phased genomes for all species were generated using hifiasm v0.19.9 (Cheng *et al*., 2021) using 20 threads and default parameters. For each species, three assembly files were output and used for downstream analyses: (i) the phased consensus contig graph of haplotype 1, (ii) the phased consensus contig graph of haplotype 2, and (iii) the raw unitig GFA file containing all possible haplotype information. Raw unitig GFA files are termed ‘raw’ because hifiasm has not performed bubble popping and duplicate purging on this GFA version. Bubble popping is the hifiasm algorithm that removes or selects one unitig at small bubbles due to their high probability of arising from sequence error or *de novo* mutation. Purging removes duplicate assembly regions (Cheng *et al*., 2021).

After conversion from GFA files to FASTA format using gfa2fa from gfatools v0.5 (Li), AFPs were annotated in the haplotype-phased assemblies for each species in order to estimate AFP III copy numbers that were later evaluated for accuracy. AFP III annotation was also performed in raw unitig graphs in FASTA format to provide putative copy numbers for each unitig. These were later used to evaluate the effect of assembly uncertainty on CNVs. Annotations of AFP III genes were performed using Exonerate v2.4.0 with a query sequence of the translated *Zoarces americanus* AFP set to the parameter ‘protein2genome’ (Slater and Birney, 2005) and a maximum intron size of 10 kb (Bogan *et al*., 2024). Intact AFP genes were differentiated from all Exonerate hits using three criteria: (1) the presence of start codon, (2) the presence of AFP III exons 1 and 2 absent large deletions, and (3) the absence of a premature stop codon. The number of functional AFP gene copies was counted for each contig in the phased contig graphs of haplotypes 1 and 2 and each unitig in the raw unitig GFA.

In order to verify the accuracy of Exonerate across substantial variation in AFP sequences between species and isoforms, Exonerate hits from the *Z. americanus* query in each species were compared with hits from an Exonerate query using an AFP amino acid consensus sequence from the target species. These consensus sequences were generated from a FASTA file containing all AFP hits output by Exonerate using the *Z. americanus* query sequence. The consensus sequence was generated from a multiple sequence alignment of these AFP nucleotide sequences, performed using MAFFT v7.526 with the FFT-NS-2 method (Katoh *et al*., 2002). Counts of intact AFP genes were compared between the *Z. americanus* and consensus sequence queries of the phased contig graphs of haplotypes 1 and 2.

In addition to the ancestral AFP arrays at which Zoarcoidei AFP arose, the genomes of *Anarhichas lupus*, *Anarhichas minor*, *Leptoclinus maculatus*, and *Zoarces americanus* also contain translocated AFP arrays (Bogan *et al*., 2024) that were analyzed. Ancestral arrays were distinguished from translocated arrays by the presence of upstream and downstream genes that exhibit conserved synteny and flank the ends of the ancestral array (Deng *et al*., 2010; Hotaling *et al*., 2023). Annotation of the upstream *sncgb* gene in the haplotype-phased assemblies and unitig GFA was performed using Exonerate v2.4.0 with a query *Gasterosteus aculeatus sncgb* protein sequence set to the parameter ‘protein2genome’ (Slater and Birney, 2005). The same annotation was performed with a query protein sequence of the *G. aculetaus ldg3b* gene, which is downstream of the ancestral array in Zoarcoidei.

### 2.3. Detecting genome misassembly and effects on inferred copy number

The effects of genome assembly uncertainty on measurement of AFP CNVs were first quantified by analyzing the raw unitig GFA in Bandage v0.8.1 (Wick *et al*., 2015). The raw unitig GFA contains information on all possible assemblies of unitigs to generate a given contig. For each AFP array, every possible contig that could be produced from the GFA was determined by calculating all directed, acyclic paths that connect the 5’-most unitig to the 3’-most unitig of a unitig network. Large bubbles in the raw GFA representing heterozygous sites corresponded to haplotype bubbles in the purged GFA after undergoing bubble popping and purging. At these heterozygous sites, small bubbles represented assembly uncertainty of a given haplotype. Therefore, analysis of raw GFA files not only enabled extraction of all possible contig assemblies for an unphased assembly, but also for haplotype-phased regions. The AFP copy numbers of each unitig were previously annotated and the copy number of each possible haplotype contig was determined by summing the copy number of all unitigs included within a given contig iteration (one of *n* possible contigs generated from a GFA). A margin of error was quantified for each haplotype by determining the minimum and maximum AFP copy number between all possible contigs, resulting in a measurement of uncertainty in haplotype copy due to misassembly.

In regions where assembly of one haplotype could not be reasonably distinguished from the other due to interconnected assembly paths through a possible haplotype bubble (Fig. 1, misassembly), the same margin of error was applied to both haplotypes—representing the minimum and maximum AFP copy number for all paths through the AFP region. In order to standardize the variance in copy number estimates between species with different AFP copy numbers, a percent uncertainty in copy number was calculated for each haplotype by dividing the margin of error by the maximum copy number. A Bayesian regression (Bürkner, 2017; Makowski *et al*., 2019) of the relationship between percent uncertainty and median AFP copy number of arrays was used to calculate the rate of change in uncertainty with increasing array copy number (see Supplemental Methods).

This workflow was automated in the script gfa_parse.py (https://github.com/snbogan/gfa_parser). The script reads in six arguments: a raw unitig GFA file, two lists of unitigs phased to each haplotype in a contig of interest, ‘start’ and ‘end’ unitigs flanking the 5’ and 3’ ends of each haplotype, and an output name (Fig. 3A). The script identifies all possible contiguous paths through the flanking unitigs of each haplotype that are directed and acyclic using the networkx package (Hagberg *et al*., 2008). It also creates an index of unitig names and their associated sequences. The directed, acyclic paths ensure that (i) no assembly path loops through a unitig multiple times and (ii) that alternate unitigs within the same bubble are not included in the same path. The script then extracts the FASTA sequence of each unitig and, for a given path, concatenates the FASTA sequences in the order of the assembly path while accounting for overlapping unitig sequences and strandedness of overlaps. *gfa_parser* can be run in phased mode or unphased mode to resolve uncertainty in regions of a haplotype-phased assembly or primary assembly (see Supplemental Methods). This process is repeated for all possible assemblies of each haplotype. The script exports a directory of all possible contigs for both haplotypes in FASTA format. To quantify the influence of assembly uncertainty on AFP copy number detection, AFP genes were annotated across *n* possible contigs of both haplotypes of each species using Exonerate as described above. We developed settings for *gfa_parser* for use on GFA files output by Verkko (Rautiainen *et al*., 2023), minigraph (Li *et al*., 2020), and Shasta (Shafin *et al*., 2020) in addition to hifiasm (see Supplemental Methods). In Supplemental Methods, we describe an application of *gfa_parser*’s approach for measuring genome-wide assembly uncertainty using a script called gw_paths.py (Fig. S5).

**Figure 3.**
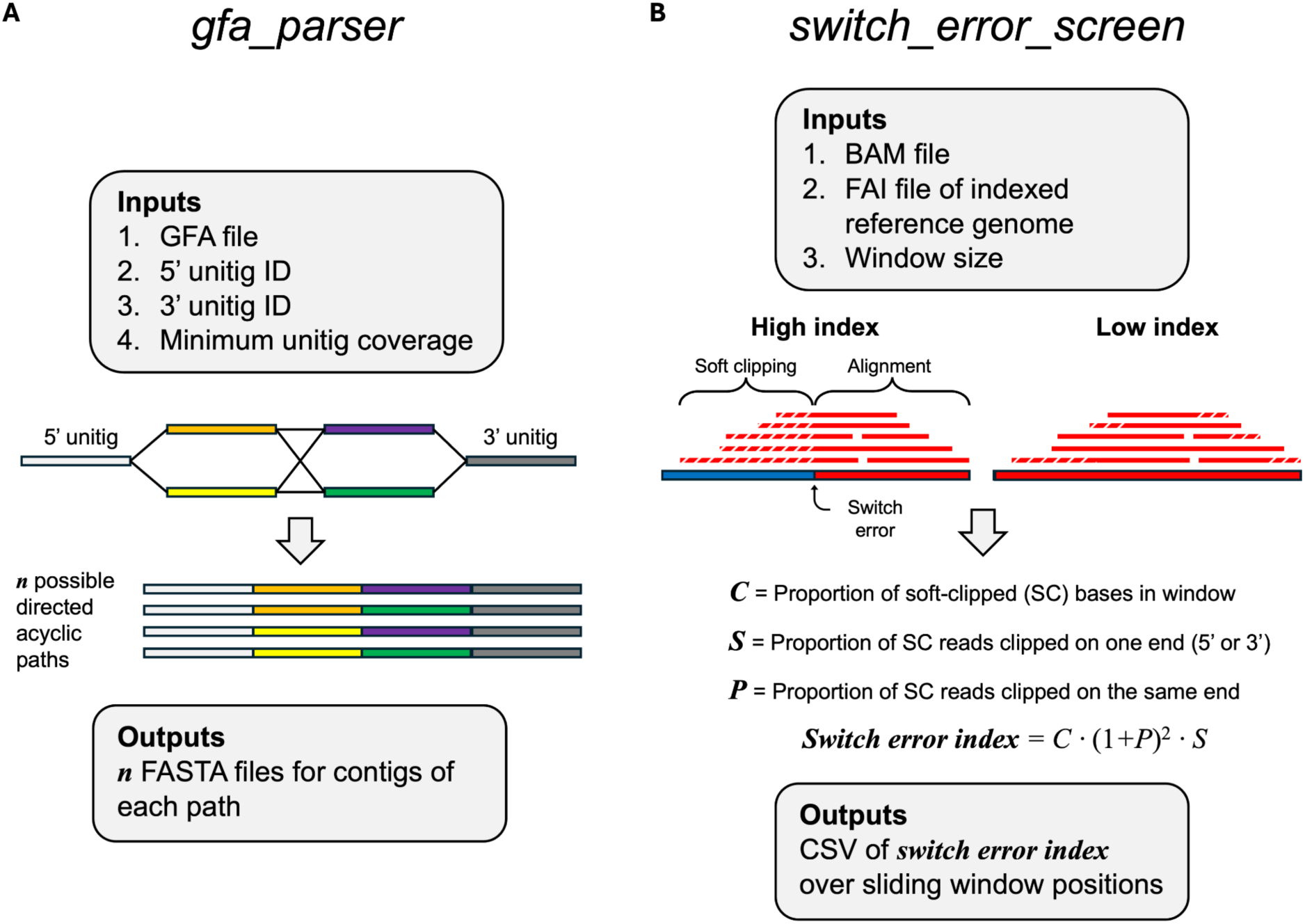
Diagram of bioinformatics tools: gfa_parser and switch_error_screen. (A) Graphical representation of *gfa_parser*, which outputs FASTA sequences for all possible paths through a defined section of a graphical file assembly (GFA) file that are (i) directed, multiple alternative unitigs in the same bubble are not included in the same path, and (ii) acyclic, no assembly path loops through a unitig multiple times. (B) Illustration describing *switch_error_screen,* which calculates a switch error index value across a sliding window based on the proportion of soft clipping on either the 5’ or 3’ end across multiple reads. Two haplotypes are represented by color (red vs. blue). Soft clipping is represented by diagonal white lines through aligned reads.

### 2.4. Screening for switch errors

Switch errors were identified by re-aligning haplotype-phased reads and unphased reads to the phased assembly FASTAs. Alignment was performed using pbmm2 v1.14.99 (/github.com/PacificBiosciences/pbmm2) with default parameters, a wrapper of minimap2 (Li, 2018) optimized for alignment of HiFi reads. Phasing information was extracted from hifiasm log files. BAM files were visualized alongside the haplotype 1 and 2 assemblies and the previously generated Exonerate AFP annotations in Integrative Genomics Viewer (IGV) v2.18.4 (Robinson *et al*., 2011). Soft-clipping was visualized in IGV and used as an indicator of switch error under the expectation that improper phasing in a haplotype bubble would result in soft clipping of reads at one side of a switch error junction.

We developed a script to automatically identify regions of high switch error likelihood and flag regions for further inspection. This script (*switch_error_screen*) is available on Github (https://github.com/snbogan/switch_error_screen). The script reads in a BAM alignment of all hapmers and unphased reads against a phased genome FASTA file. Over a 10 kb sliding window, the script calculates an index of switch error by measuring three variables: the proportion of soft clipped reads within the window (*C*), the skewness of soft clipping toward 5’ and 3’ ends among these reads (*S*), and polarization of soft-clipping skewness (*P*) as shown in Figure 3B. Each of these metrics are extracted from soft clipping data retained within the BAM CIGAR string. *C* was calculated as the number of soft clipped bases divided by the sum of all matched, inserted, and deleted bases. *S* was calculated as the variance in the per-read difference between the number of left and right soft-clipped bases. *P* was determined by the absolute difference between the number of reads clipped on the left versus the right, normalized by the total number of clipped reads.

Under switch error caused by incorrect phasing or misassembly, we expect to observe junctions at which raw reads exhibit (i) a high proportion of soft clipping (*C*), (ii) soft clipping on one side of each affected read (*S*), and (iii) soft clipping on the same side of a switch error junction among affected reads (*P*). The script then calculates the index of switch error risk according to Equation 1.

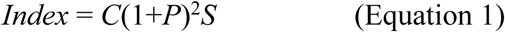

Here, (1+*P*)^2^ ensures that the index increases more rapidly as polarization deviates from zero. When polarization is zero (P=0), the term evaluates to 1 rather than 0, and does not eliminate risk associated with skewed soft-clipping being evident within a region.

The sensitivity of *swtich_error_screen* to detect patterns of soft-clipping consistent with switch errors was evaluated by applying the the script to (i) a FASTA containing a simulated switch error in the *P. gunnellus* AFP array and (ii) realignments of HiFi reads back to their corresponding reference genomes. The FASTA containing a simulated switch error was produced by first performing a chromosome alignment of *P. gunnellus* AFP array haplotypes 1 and 2 using Mauve (Darling *et al*., 2004). FASTA sequences from the two haplotypes were trimmed and concatenated such that the 5’ end of haplotype 1 and the 3’ end of haplotype 2 fused at the start of an extended region at which the two haplotypes shared no sequence homology (Fig. S1). This imposed an artificial switch from haplotype 1 to 2 at a region where the two alleles diverged.

## 3. Results

### 3.1. Validation and performance of gfa_parser

The performance of *gfa_parser* was evaluated by measuring concordance between (i) contig FASTA sequences produced by the script when input with raw hifiasm GFA files and (ii) contigs that were assembled through manual inspection and concatenation of unitig sequences in raw hifiasm GFA files. In each AFP array for which haplotypes could be resolved, *gfa_parser* computed the same number of directed acyclic paths through haplotype unitigs that was calculated during manual inspection (Fig. 3A). Contigs output by *gfa_parser* shared the same length and AFP copy number as manually assembled contigs. The functionality of *gfa_parser* was also tested on GFA files generated by the genome assemblers Verkko (Rautiainen *et al*., 2023), Shasta (Shafin *et al*., 2020), and the pangenome graph assembler minigraph (Li *et al*., 2020). *gfa_parser* output the same number of contig FASTA files as directed acyclic paths calculated through manual inspection. These results confirmed the efficacy of *gfa_parser* for automatically extracting multiple contig assemblies from multiple GFA assemblers.

### 3.2. Misassembly in antifreeze protein tandem arrays

Sequencing of *L. dearborni* achieved a combined yield of 46.20 gigabases of HiFi data, a mean read length of 8.98 kb, and a read N50 of 11.96 kb. Sequencing of *Z. americanus* achieved a yield of 19.79 gigabases of HiFi data, a mean read length of 11.34 kb, and a read N50 of 9.30 kb. Based on genome sizes of the primary unphased assemblies, these yields achieved coverages of 63.94x for *L. dearborni* and 31.69x for *Z. americanus*. Contig N50s were calculated for the primary unphased assemblies as 7.08 Mb for *L. dearborni* and 3.22 Mb for *Z. americanus*.

To understand the effects of genome misassembly on measurement of AFP CNVs, variance in copy number between different paths through the graphical fragment assembly (GFA) was assessed. Because of the highly repetitive structure of AFP tandem arrays, several species exhibited multiple possible assemblies through unitigs spanning the array. Assembly uncertainty was also evident in haplotypes of heterozygous regions: large GFA bubbles corresponding to distinct haplotypes branched into assemblies that contained small bubbles, which demonstrated uncertainty within haplotype assemblies (Fig. 4A). This created measurable uncertainty in haplotype-specific copy numbers in some species (Fig. 4B).

**Figure 4.**
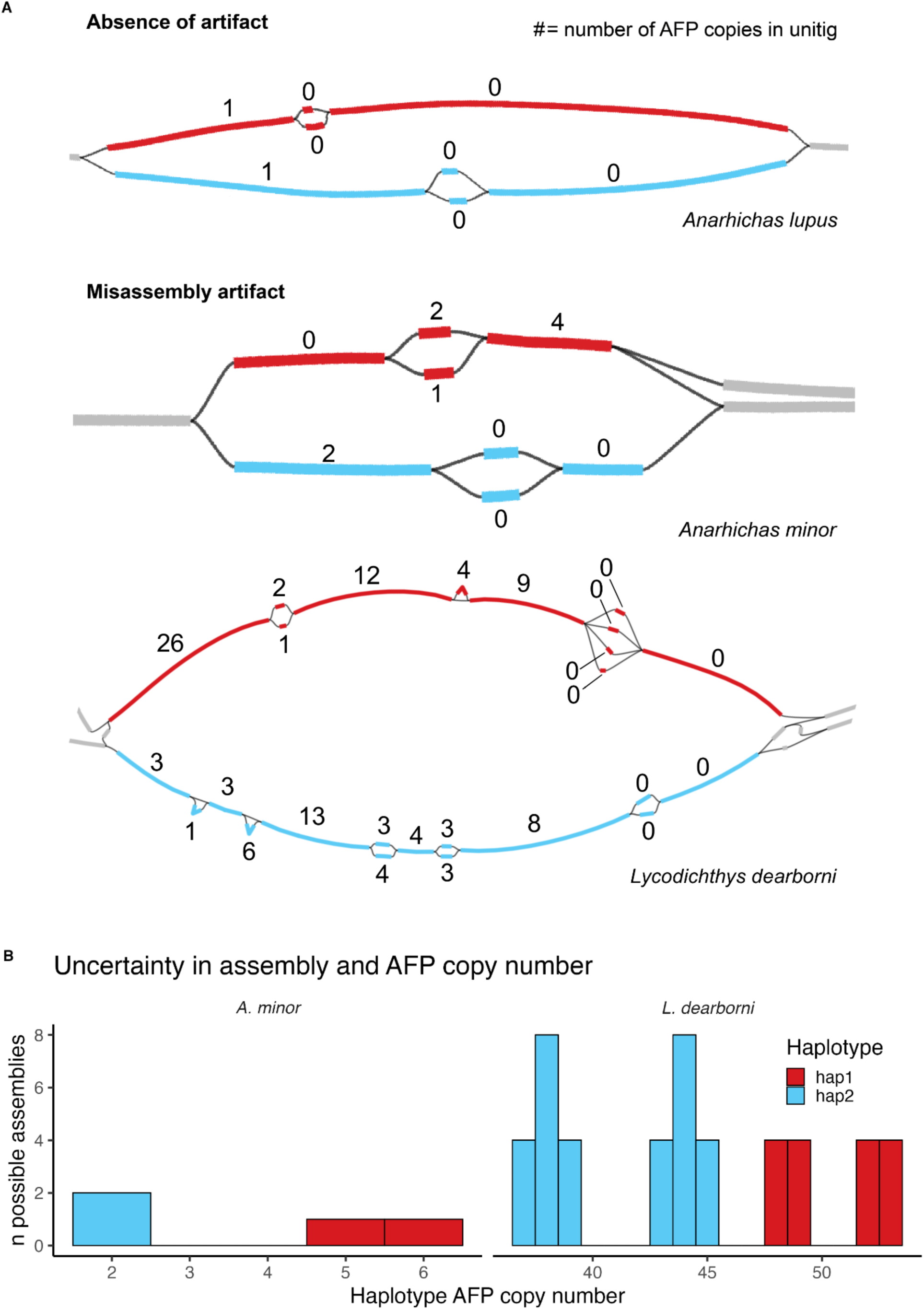
Effect of assembly uncertainty in antifreeze protein tandem arrays on copy number detection. (A) Assembly string graphs of raw unitig GFA files are shown for *Anarhichas lupus*, *A. minor*, and *Lycodichthys dearborni* that span the ancestral AFP array of each species. Thick bars represent unitig nodes. Thin black lines represent edges, which denote partial overlaps between unitig ends. Haplotype bubbles are differentiated by color: haplotype 1 = red; haplotype 2 = blue. (B) Variation in AFP copy number within haplotypes is plotted across 2 possible assembly contigs of *A. minor* haplotypes 1 and 2 for the ancestral AFP array and 16 and 36 possible assembly contigs in haplotypes 1 and 2 of the *Lycodichthys dearborni* ancestral array. A histogram is used to visualize the number of assembly iterations annotated with a given AFP copy number. Assembly iterations were parsed from the raw hifiasm GFA file using *gfa_parser* (see Methods). AFP CNVs among possible contigs demonstrated differences of at least 2 copies between *A. minor* haplotypes and at least 3 copies between *L. dearborni* haplotypes.

The number of possible assemblies that could be generated from raw GFAs at AFP arrays ranged from one possible assembly per haplotype in the *P. gunnellus* ancestral array to hundreds per haplotype in a translocated array of *L. maculatus*. A large bubble existed at a translocated array of *L. maculatus* in the raw GFA, but did not possess two clear sets of assembly paths consistent with distinct haplotypes and, therefore, the assembly could not reliably resolve possible haplotype assemblies. Examples of GFAs that could resolve possible haplotype assemblies are provided in Figure 4A. The GFA of the unresolved *L. maculatus* AFP array is visualized in Figure S2. Similarly, haplotype assemblies could not be resolved for the *Z. americanus* and *A. lupus* ancestral arrays.

In both unresolved and resolved cases, misassembly artifacts affected the estimation of phased AFP counts in resulting haplotype contigs because the hifiasm algorithm chooses one of many pathways through the unitigs for each haplotype contig, potentially misrepresenting assembly or copy number if assembly uncertainty is not taken into account. Figure 4A contrasts a raw GFA from *A. lupus*, which shows no variation in AFP copy number between possible assembly paths, with raw GFAs from both *A. minor* and *L. dearborni,* where multiple assembly paths with different AFP copy number can be derived from one contig. Here, *A. minor* haplotype 1 had a measurable uncertainty in AFP copy number of 1 copy. *L. dearborni* haplotype 1 displayed an uncertainty in AFP copy number of 5 copies, while haplotype 2 had an uncertainty of 8 copies. Note that while the two haplotypes share a start and end unitig, the number and order of AFPs across unitigs is distinct between either side of the haplotype bubble due to substantial differences in the locations of AFP genes between haplotypes. In regions where misassembly artifacts create uncertainty in AFP CNVs, uncertainty can be quantified as (i) difference between the minimum and maximum possible copy numbers for each haplotype or (ii) the distribution of possible AFP copy numbers per haplotype. Examples of each metric for AFP CNVs uncertainty are present in Figure 4B where distributions of uncertainty and minimum and maximum copy numbers are plotted.

The number of AFP copies varied largely between species, and it was expected that the percentage of uncertainty in AFP copy number might increase in repetitive, high-copy number arrays. However, percent uncertainty in AFP copy number did not correlate with mean AFP copy number. The slope of median AFP copy number’s effect on percent uncertainty was estimated to be 0.0 with a 95% confidence interval of −0.02 – 0.02 (Table 1; Fig. S3). *L. maculatus* showed both the highest copy number at a single translocated array and the greatest uncertainty in copy number, with a 64 copy difference between the assembly pathways of minimum and maximum AFP copy number (57% uncertainty). However, the second highest uncertainty in copy number was in the ancestral array of *A. lupus,* where 6 of 11 copies (55%) were not present in all possible assemblies. In both cases, the GFA contained pathways connecting both sides of the heterozygous bubble, preventing distinct haplotypes from being resolved during analysis. All other species exhibited a percent uncertainty in AFP copy number in at least one array, except for *P. gunnellus,* with uncertainty ranging from 9% to 33%. Measurements of uncertainty in copy number were recapitulated between manual inspections of GFA files and automated analysis using *gfa_parser*.

**Table 1.**
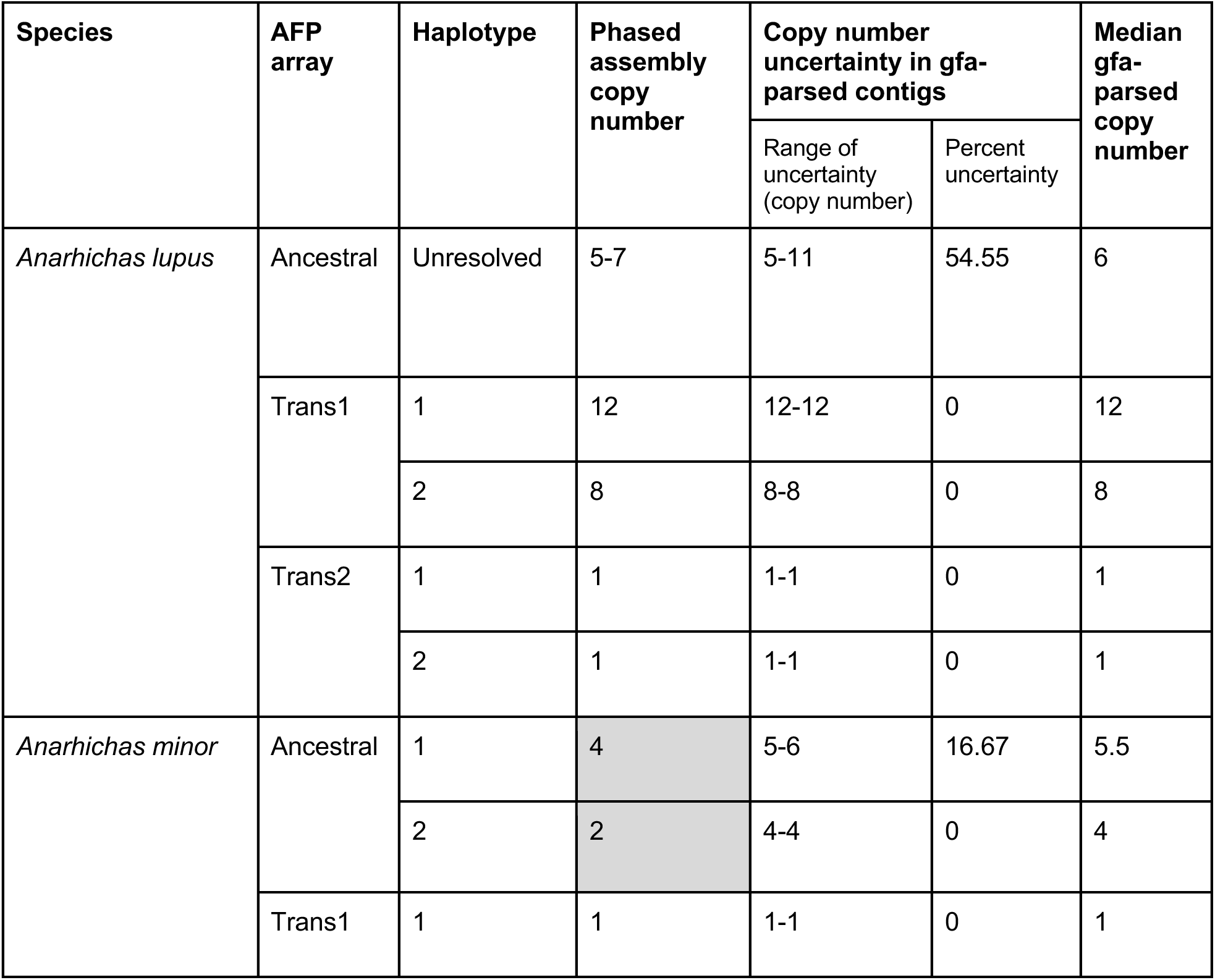
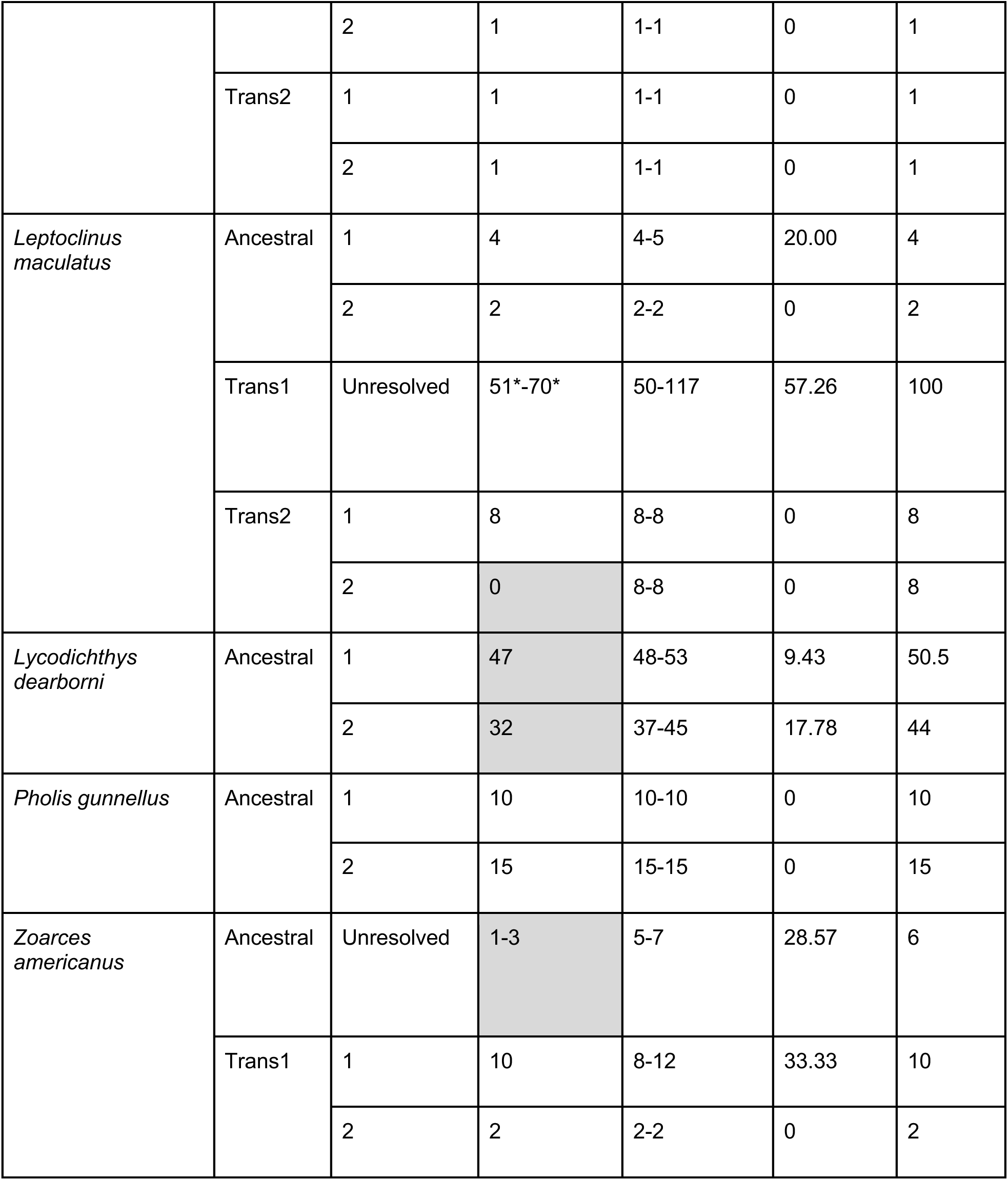
Impact of misassembly artifacts on AFP CNVs in haplotypes across species. Effects of assembly uncertainty are shown across species with functional AFPs in ancestral AFP arrays and arrays translocated out of the ancestral array to different chromosomes (’Trans’). Number of AFP copies estimated through annotation of haplotype-phased assemblies is contrasted with the median number of copies determined during analysis of the raw unitig string graph. Copies deemed uncertain are represented as both a number of copies between minimum and maximum AFP counts and a percentage of maximum copies in that haplotype. Grey boxes represent instances where AFP copy numbers of final phased contigs produced by hifiasm were lower than the range of possible haplotype copy numbers inferred from raw graphical fragment assembly (GFA) files. * = a contig of the raw GFA that was split into multiple contigs of final phased assemblies.

In 7 of 25 AFP array haplotypes, AFP copy numbers of final phased assemblies were lower than the minimum possible copy number inferred from raw GFA files. This proportion was 6 of 23 when collapsing unresolved haplotypes into single observations as shown in Table 1. In these cases, AFP copy numbers of final phased contigs were 1-20 copies lower than the range of possible haplotype copy numbers inferred from GFAs (Table 1). Final or ‘primary’ contigs produced by hifiasm are subject to bubble popping and duplicate purging and represent a single assembly path of a purged GFA file (Cheng *et al*., 2021). Alignment of gfa-parsed contig iterations against final phased contigs and inspection of GFA metadata revealed that the downward bias was attributed to removal of small bubbles from the purged GFA that contained unitigs with AFP copies. In raw GFA files, unitigs within these small bubbles had low read coverage of 1-2 reads per unitig (Fig. S4). While hifiasm bubble popping generally selects one unitig of highest read support from all possible unitigs in a small bubble, the algorithm will remove all unitigs and purge bubbles when read support is low for each unitig (Cheng *et al*., 2021).

### 3.3. Switch errors were unapparent in hifiasm haplotype assemblies

Switch errors were screened for by scanning for abundant soft clipping of phased reads aligned to their corresponding haplotype assembly, with soft clipping occurring on one side of a junction demonstrating a switch in haplotype across a phased assembly (Fig. 3B). Screening was performed using manual inspection of alignments and automated bioinformatic analysis. Realignment of HiFi reads to the simulated switch error FASTA sequence resulted in several positions at which switch errors caused polarized soft-clipping and high *switch_error_screen* scores (Fig. 5). A large peak in switch error risk is evident at the artificial switch error visualized in Figure 5. This peak is attributed to a region with polarized soft clipping on the 5’ and 3’ ends of a sequence that is homologous between haplotypes (Fig. 5). *switch_error_screen* also demonstrated sensitivity to detecting smaller levels of polarized soft clipping within unaltered contigs. These results confirmed the sensitivity of the tool to detect soft clipping signatures of switch errors.

**Figure 5.**
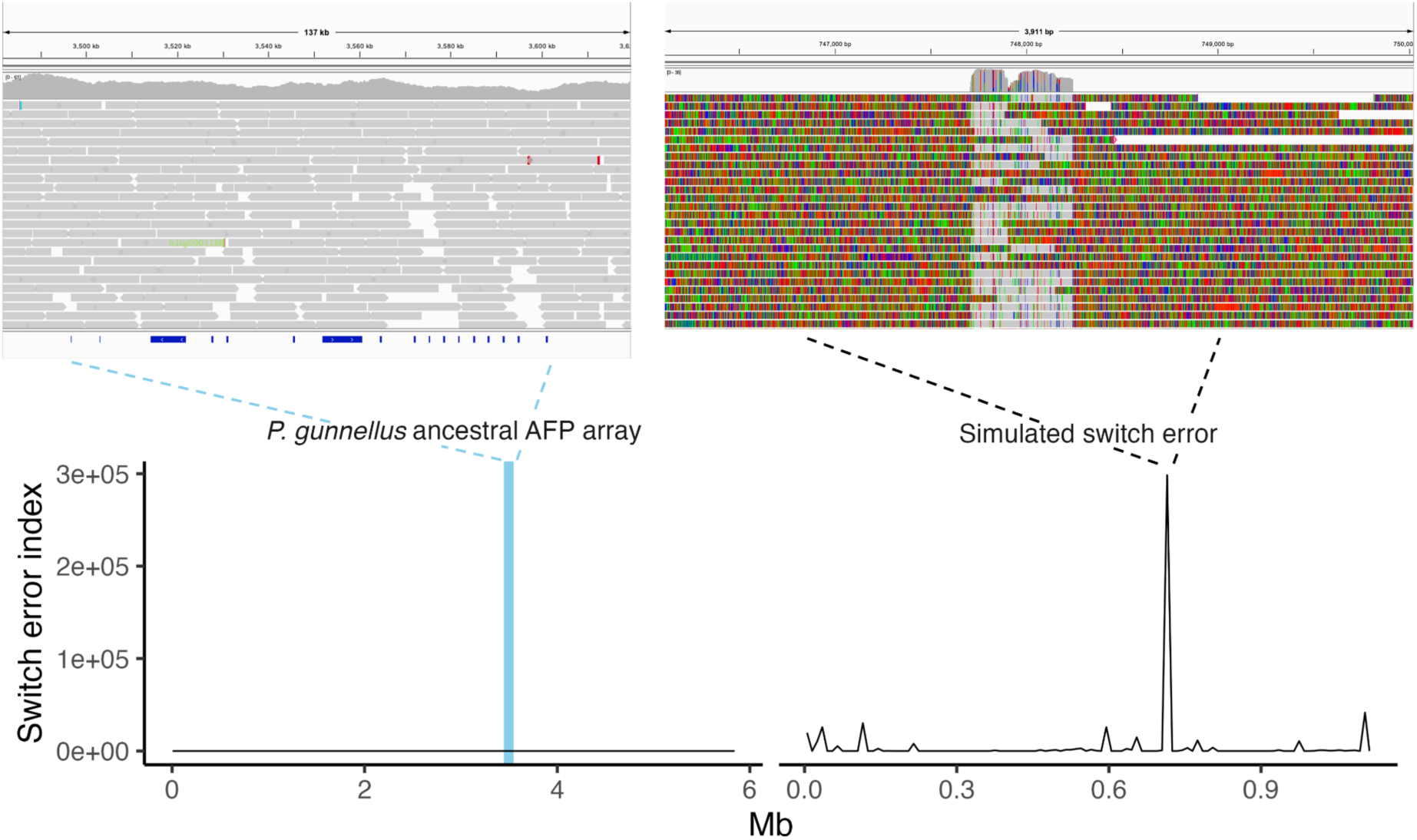
Switch errors in antifreeze protein tandem arrays. Realignments of HiFi reads back to reference genome species for *Pholis gunnellus* (left) and simulated switch errors (right) are visualized in IGV (top row) and input to the *switch_error_screen* tool, which calculates the risk of a switch error according to soft clipping (bottom row). An artificial switch error was introduced to the ancestral antifreeze protein array in the *P. gunnellus* reference genome for the purpose of comparison to an unaltered *P. gunnellus* array with low switch error risk. Soft clipped bases are colored in the IGV visualizations. Ranges of AFP gene annotations are visualized on the bottoms of IGV images as dark blue rectangles. A blue highlight in the left switch error index plot denotes the range of ancestral AFP tandem arrays.

While soft clipping was observed across phased assemblies, soft clipping consistent with switch errors were not apparent in Zoarcoidei AFP arrays. Figure 5 displays the results of *switch_error_screen* in a typical phased assembly versus a simulated phasing switch error, where abundant soft clipping in a tandem array of AFPs suggests errors when phasing reads to the correct haplotype. In this case, multiple switches from one haplotype to another in the middle of a contig could affect inferred haplotype AFP copy number by switching the haplotype assemblies on the soft-clipped side of the switch error. However, the hifiasm phased assemblies that we report here all appeared to be robust against switch errors.

### 3.4. Haplotype diversity in copy number of antifreeze protein genes in polar fishes

Haplotype-specific AFP copy number was compared within and across Zoarcoidei possessing intact AFP genes after controlling for the effects of misassembly on inferred copy number. AFP CNVs between haplotypes were present in all species and over half of all AFP arrays for which haplotypes were resolved. In certain instances, such as translocated arrays of *A. minor*, assembly artifacts caused no uncertainty in AFP copy number and no CNVs between haplotypes were observed. In *P. gunnellus*, assembly artifacts did not induce uncertainty in copy number and haplotype-specific CNVs were present (Fig. 6, Table 1). In some cases, the presence of artifacts created variable estimations of copy number, but CNVs between haplotypes still occurred. For example, the translocated array of *Z. americanus* had an uncertainty of four copies in one haplotype due to misassembly, yet a minimum difference of six copies between haplotypes remained. In three arrays, misassembly artifacts created too much uncertainty to resolve distinct CNVs between haplotypes. In a translocated *L. maculatus* array, for example, the effects of misassembly prevented estimations of differing copy numbers between haplotypes (Fig. 6, Table 1).

**Figure 6.**
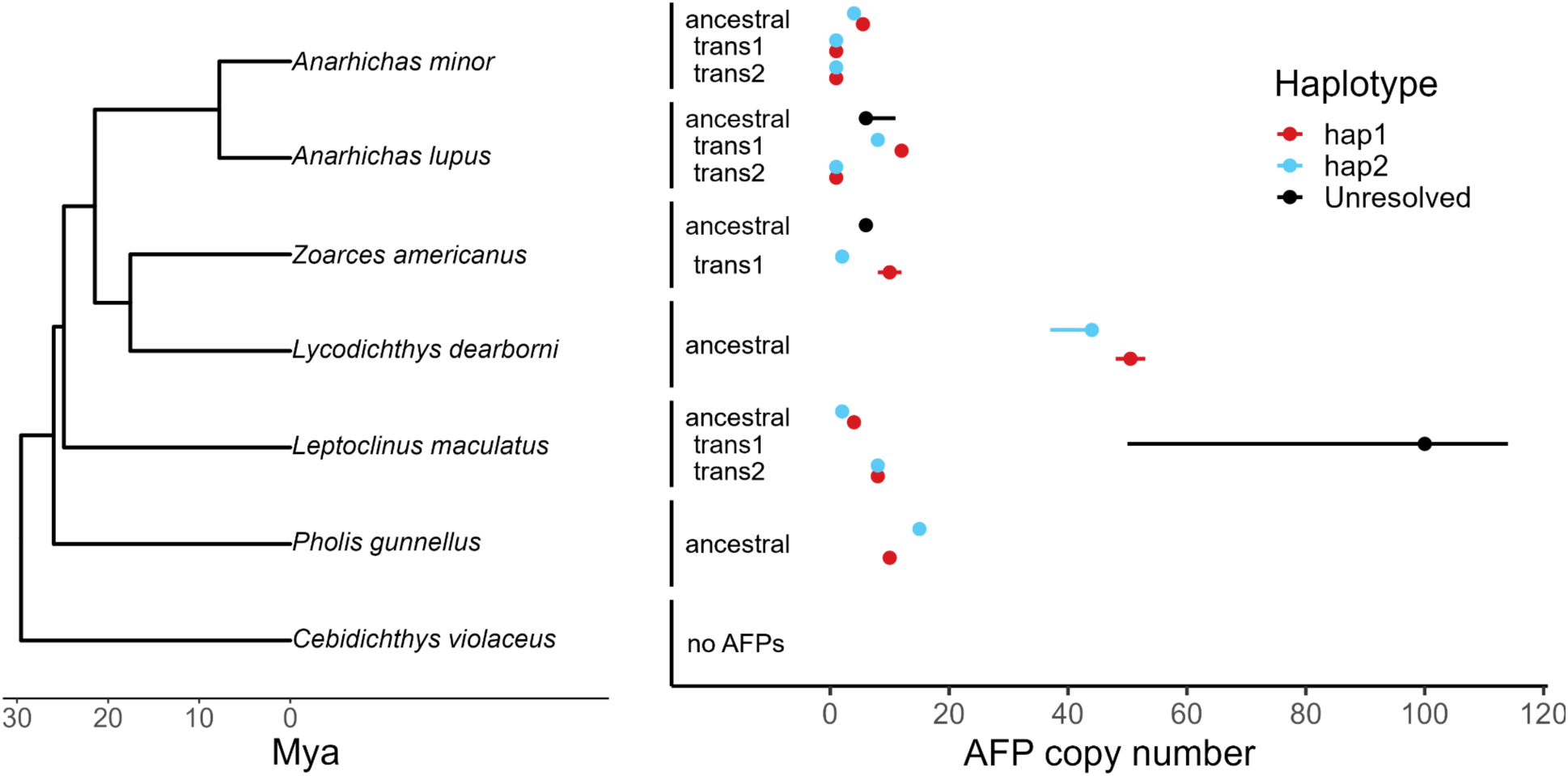
Haplotype diversity in antifreeze protein gene copy number across species. Time-calibrated species tree of Zoarcoidei was generated from a multilocus alignment (Rabosky *et al*., 2018) of all six species with intact AFP genes and a zoarcoid outgroup lacking AFP. Numbers of AFP copies per haplotype are listed for ancestral and translocated arrays. Points represent the median AFP count from all possible assemblies of the AFP array inferred from raw graphical fragment assemblies (GFA files). Error bars represent the minimum and maximum copy number across possible assemblies of an array. Within-species haplotype diversity in AFP copy number is represented by pairs of points and error bars that are nonoverlapping, demonstrating differences in AFP copy number after controlling for assembly uncertainty.

Overall, CNVs between haplotypes were observed in all six species with functional AFPs and 6/10 haplotype-resolved AFP arrays (Fig. 6). Of the remaining seven arrays with no CNVs between haplotypes, four showed no evidence of variation, while the other three contained assembly artifacts that created too much uncertainty to resolve distinct copy numbers between haplotypes (Table 1). In species with more than one AFP array in their genome, CNV was observed between haplotypes only in one array. In general, misassembly errors created the greatest impact on estimations of copy number. Additional analyses of switch error did not result in the discovery of errors that affected copy number counts (Table 1).

## 4. Discussion

Haplotype-aware approaches are advancing genomic research in natural populations (Leitwein *et al*., 2020; Secomandi *et al*., 2025), particularly investigations of large structural genomic variation (SV). For example, phased genomes and pangenomes enable the detection and phasing for SVs of greater length than detection via mapping of long reads (Ebert *et al*., 2021). In samples that lack parent-offspring trio data traditionally used for haplotype phasing (Marchini *et al*., 2006), assembly errors and switch errors in phased genomes have the potential to introduce error in haplotype-specific SV detection even when phasing with Hi-C sequences is performed (Cheng *et al*., 2022; Li and Durbin, 2024). We have described new methods for (i) measuring assembly uncertainty in haplotype-phased genomes, and (ii) detecting switch errors caused by phasing or misassembly (Fig. 3). We applied these tools to pseudo-phased hifiasm *de novo* assemblies of polar fishes in the suborder Zoarcoidei, which exhibit a rapidly evolving tandem array of antifreeze protein (AFP) genes (Hotaling *et al*., 2023; Bogan *et al*., 2024). After applying these tools, we report the first genomic analysis of intraspecific diversity in AFP genes and haplotypes among polar fishes.

Assembly uncertainty was abundant in repetitive regions of phased genomes and introduced error to measurements of haplotype AFP copy number in almost all arrays (Table 1; Fig. 4).We found no evidence of switch errors attributed to improper phasing by hifiasm and demonstrated the sensitivity of our method for detecting phasing switch errors using simulation (Fig. 5). After quantifying and accounting for uncertainty caused by these artifacts, we observed significant haplotype diversity in AFP copy number in 6 out of 13 sequenced AFP arrays and in each Zoarcoid species exhibiting intact AFP genes (Fig. 6). Our results provide a workflow for rigorously evaluating large haplotype-specific SVs in phased genomes and pangenomes and demonstrate that intraspecific variation and haplotype diversity in AFP copy number are pervasive across Zoarcoidei. Below, we discuss how our findings inform current practices in pangenomics and the evolution of AFPs in polar fishes.

### 4.1. Novel tools for quantifying assembly uncertainty

Hifiasm is a popular tool for *de novo* assembly of phased genomes using long-read PacBio HiFi data. Hifiasm also produces valuable metadata for inferring and quantifying assembly uncertainty in its primary and phased genomes. These metadata are contained in the raw unitig graphical fragment assembly file or ‘r_utg.gfa’ (Cheng *et al*., 2021). Similarly to hifiasm, Verkko (Rautiainen *et al*., 2023) and Shasta (Shafin *et al*., 2020) can output intermediate GFAs that contain assembly graphs with all possible nodes and edges. While older versions of the assembler Canu (Koren *et al*., 2017) output a contig assembly graph, newer versions do not output a GFA due to incompatibility with the GFA2 format. Although GFA files are commonly output by long-read assemblers, some assemblers that use short reads, such as SPAdes (Prjibelski *et al*., 2020) also output an assembly graph in GFA format. Inspecting GFA files for signatures of assembly uncertainty or misassembly is commonly done using manual, visual inspection in graphical GFA viewers such as Bandage (Wick *et al*., 2015) and GfaViz (Gonnella *et al*., 2018).

Here we described and applied a novel tool named *gfa_parser*, a python script that exports FASTA sequences representing all possible contiguous assemblies of a region contained within a GFA (Fig. 3A). The script reads in a GFA and lists of unitigs spanning two haplotypes or a range between two unitigs. Using network analysis, the script constructs all possible directed acyclic paths through unitigs of the region or its two haplotypes. It then concatenates FASTA sequences of unitigs in their directed order across each assembly path. While we applied *gfa_parser* to a specific case study, the tool can be used with a GFA file for any system produced by any assembly algorithm to extract the full range of contiguous FASTAs that are possible at a given region. For data without phasing information, the unphased mode of *gfa_parser* can be used. Here, *gfa_parser* was used to determine whether gene copy number expansion/contraction in a species or lineage is attributed to misassembly by measuring gene copy number across contig iterations produced by the script. However, the contig iterations can be used to determine the effects of misassembly on detection of large SVs, synteny, repetitive element content, or any application that uses a reference genome FASTA. As input, *gfa_parser can read in* phased, unphased, short-read, or long-read assemblies in GFA format.

Leveraging phased genomes to measure haplotype diversity within individuals or populations is increasingly common in molecular evolution and has been used to measure haplotype diversity in CNVs (Chen *et al*., 2022; Frandsen *et al*., 2023) and other structural variants (Schwessinger *et al*., 2018; Hoopes *et al*., 2022; Takeuchi *et al*., 2022). However, these studies were unable to examine whether haplotype diversity was attributed to misassembly, largely because bioinformatic tools were not available to facilitate such a test. Evaluating haplotype diversity against the misassembly of phased genomes is critical because likelihoods of missassembly and haplotype diversity increase in repetitive regions where rates of CNVs and SV are high (Tørresen *et al*., 2019). In Zoarcoidei, we found that misassembly and CNVs are pervasive in repetitive AFP arrays.

New technologies show potential for resolving repetitive arrays. Recent studies in human genetics have used optical genome mapping to resolve complex structural variation in the repetitive amylase locus (Yilmaz *et al*., 2024). Optical mapping has also helped characterize complex cancers by analysis of chromosomal rearrangements, deletions, and gene fusions (Vieler *et al*., 2023). Optical genome mapping holds promise for resolving SVs in repetitive regions and is optimized to detect SVs down to a resolution of 500 basepairs, a finer resolution than the average length of the Zoarcoidei AFP gene. Our results demonstrate that assembly uncertainty in repetitive arrays of AFP genes is likely ubiquitous across Zoarcoidei species. Therefore, optical mapping may improve the resolution of structural variants in this region and tandem gene arrays of other systems.

Another solution to resolve these repetitive arrays includes increased read length and sequencing depth. New Oxford Nanopore Technologies (ONT) sequencing methods generate ultralong reads that can reach over a megabase and could span the entire AFP array in a single read. Genome assemblers such as Verkko perform integrative contig assembly of both HiFi long-read data and ONT ultra-long reads (Rautiainen *et al*., 2023). Assembly uncertainty should decrease in complex regions using assembly with ultralong reads, and we propose that successful reductions in uncertainty can be validated using *gfa_parser* and gw_paths.py.

Included in the *gfa_parser* repository, the gw_paths.py script enables quantification of assembly uncertainty in GFA files at a whole-genome scale. Efficiently calculating, reporting, and analyzing multiple paths through GFAs is an area of need in genome assembly and pangenomics (Sohn and Nam, 2018; Baaijens *et al*., 2022; Sirén and Paten, 2022; Secomandi *et al*., 2025). gw_paths.py offers a unique tool for evaluating the quality of GFA files during genome assembly and measuring the structural diversity of GFA paths.

*switch_error_screen* automates the detection of switch errors in phased genomes or pangenomic graphs. The approach taken by *switch_error_screen* requires the long reads used to assemble the reference genome being evaluated. It should not be used on short read data or long reads that were not used for assembly. This is because the tool analyzes soft clipping information. The shorter a read is, the less likely it is soft-clipped during alignment. Therefore, application of *switch_error_screen* to short-read alignments risks false negatives. *switch_error_screen* is compatible with alignments of other long-read sequences such as ONT and phased assemblies of these data. However, long-read alignment algorithms such as minimap2 have popular preset parameters for HiFi and ONT genomic reads that influence the probability of soft-clipping. minimap2 ONT parameters are generally more permissive to weak alignments (Li, 2018), increasing the potential for soft-clipping and noise in the *switch_error_screen* index value.

### 4.2. Strengths and pitfalls of GFA processing for measurements of haplotype diversity

Hifiasm can produce fully-phased or pseudo-phased haploid genomes that enable analyses of haplotype diversity for a range of polymorphisms and variables (Bhat *et al*., 2021; Cheng *et al*., 2022; Engelbrecht *et al*., 2024). We evaluated the occurrence of artifacts in phased assemblies produced by hifiasm: namely, misassemblies and switch errors. We also observed biases imposed by hifiasm’s workflow that can influence measures of haplotype diversity if proper precautions are not taken. In our case study, hifiasm was robust against switch error. Of the 13 AFP arrays we studied and their respective contigs, we detected 0 instances of switch errors via analysis with our tool *switch_error_screen* and through manual inspection of realignments. These approaches quantified or looked for soft clipping of phased, HiFi long reads aligned to their respective haplotype assembly that were polarized to one side of a putative switch error junction (Fig. 3B). The categorical assignments of haplotypes in different haplotype bubbles are arbitrary and, of course, can lead to switch errors between distinct bubbles of pseudo-phased assemblies. Within haplotype bubbles, however, phasing by hifiasm appears to be robust even in highly repetitive regions.

One weakness of hifiasm’s phased haplotype assemblies is an apparent bias toward generating primary contigs that contain fewer tandem array paralogs. This bias is independent of phasing and is a consequence of the hifiasm ‘bubble-popping’ algorithm. ‘Bubble popping’ is the process hifiasm uses to filter out alternative contig assembly paths or delete short, unresolved, or low read-support regions in raw GFA files. Bubble popping is performed under the assumption that short bubbles stem from sequencing errors or somatic mutations rather than inter-haplotype variation (Cheng *et al*., 2021). However, small bubbles of low read support are common in poorly-resolved repetitive regions and are not necessarily attributed to sequencing error or somatic mutation (Leonard *et al*., 2022). Bubble popping is applied to the raw unitig GFA file (r_utg.gfa) to generate a purged GFA file (p_utg.gfa). Contiguous FASTA sequences representing the phased contigs are extracted from the purged GFA, treating alternative unitigs in the remaining bubbles as distinct haplotypes (Cheng *et al*., 2021). During bubble popping, hifiasm will completely delete small bubbles if they are associated with low-to-moderate read depth rather than selecting one unitig to incorporate in the purged GFA and resulting assembly (Cheng *et al*., 2021), effectively deleting a region that may contain paralogs of interest and fusing unitigs that share no overlap (Fig. S4).

Similarly, when constructing final phased assemblies, if multiple paths are possible through a given haplotype, hifiasm applies bubble popping to select the single best path (Cheng *et al*., 2021). This step may further bias towards assemblies that favor read depth over other variables. The long-read assembler Verkko employs a similar graph cleaning strategy based on read depth to remove alternative paths (Rautiainen *et al*., 2023). Other popular assemblers, including Shasta (Shafin *et al*., 2020) and Canu (Koren *et al*., 2017), use a best overlap strategy when multiple paths are present, choosing the read with the longest overlapping sequence with the previous read. The processing of GFA files during genome assembly is necessary for exporting high-confidence contigs, but can potentially remove biologically relevant information (Fig S4). *gfa_parser* provides a tool for evaluating the consequences of different path and unitig filtering decisions during GFA processing by outputting FASTAs of all possible contigs and their read depths. Additional metadata output by the assembler can be used alongside *gfa_parser* outputs to manually select an optimal contig. This provides flexibility for the assembly of specific, problematic regions such as repetitive tandem arrays.

In our analysis, we found that small bubbles in the raw utg.gfa contained AFP genes. Alternative unitigs of small bubbles often exhibited different non-zero AFP copy numbers. Small bubbles removed during bubble popping exhibited mean read depths of 1-3x (Fig. S4). In some cases, single unitigs removed during bubble popping contained read depths over 30x. The deletion of these regions resulted in purged GFAs and phased contigs with fewer AFP copies than were possible, according to raw GFAs (Table 1). This bias can be avoided by applying *gfa_parser* to raw utg.gfa files and/or careful inspection of the raw utg.gfa. Pangenomes and phased genomes generated using hifiasm (Leonard *et al*., 2022; Wu *et al*., 2024; Taylor *et al*., 2024) may underestimate CNVs and contig length in repetitive regions as a result of small bubble popping.

### 4.3. Haplotype diversity in copy number of antifreeze protein genes in polar fishes

The convergent evolution of antifreeze proteins in cold-adapted bacteria, plants, invertebrates, and vertebrates is a classical example of environmental adaptation (Duman, 2015; Muñoz *et al*., 2017; Wisniewski *et al*., 2020). AFPs (specifically, antifreeze glycoproteins or AFGPs) were first discovered in marine fishes of the suborder Notothenioidei (DeVries and Wohlschlag, 1969; DeVries, 1971). Research on the evolutionary genetics of AFP genes has continued to focus on fishes, where four types of AFPs and AFGPs have each independently evolved multiple times, and has almost entirely been restricted to comparative genomic approaches evaluating macroevolution across species (Ewart and Hew, 2002; Baalsrud *et al*., 2018; Zhuang *et al*., 2019; Graham *et al*., 2022; Rives *et al*., 2024). A study by Graham et al. (2022) observed intraspecific variation in AFP genes and their copy number by sequencing haplotypes of a type I AFP array in an individual diploid flounder before performing Southern blotting on multiple individuals to evaluate the frequency of these two alleles across four populations (Graham and Davies, 2024). This study provided early evidence that there is considerable intraspecific variation in AFP copy number among polar fishes, and that apparent interspecific differences in AFP genes are likely shaped in part by microevolutionary processes. Another study of AFGPs in the Atlantic Tomcod *Microgadus tomcod* used molecular cloning and fluorescence *in situ* hybridization to detect heterozygosity in an AFGP array characterized by the presence of AFGP in one allele and absence of the gene in the other. This demonstrated that haplotype diversity in AFGP and AFP may not only influence gene copy number, but gene presence/absence (Zhuang *et al*., 2018).

Our findings further support the importance of intraspecific variation in AFP evolution by demonstrating that heterozygosity and haplotype diversity in type III AFP copy number is pervasive across polar fishes of the Zoarcoidei suborder. After controlling for uncertainty in haplotype-specific copy number estimates, we detected intra-individual haplotype diversity in AFP copy number at 6 out of 13 AFP tandem arrays and in all genomes from species with intact AFP genes. This high level of intraspecific variation is likely a driver of environmental gradients in AFP copy number observed across globally-distributed Zoarcoidei species (Hotaling *et al*., 2023; Bogan *et al*., 2024) and derived multimeric AFP genes in polar Zoarcidae that likely evolved through tandem duplication (Wang *et al*., 1995; Kelley *et al*., 2010). Intra-individual haplotype diversity was evident in copy number of the ancestral AFP tandem array as well as translocated AFP arrays. We previously found that translocation-duplication contributed to AFP expansions in cold-adapted and shallow water Zoarcoidei species (Bogan *et al*., 2024). Future research should test whether high copy number haplotypes of ancestral and translocated AFP arrays increase in frequency and are positively selected for in populations inhabiting cold, shallow-water habitats. More broadly, our results suggest that haplotype-aware population genomic approaches have great potential to uncover mechanisms that shaped AFP evolution.

## Ethics Statement

Use of animal tissue samples for this research was approved by the IACUC board of the University of California, Santa Cruz under IACUC approval Kellj2302. *Zoarces americanus* was collected by the Virginia Institute of Marine Science NorthEast Area Monitoring and Assessment Program under Commonwealth of Massachusetts, Division of Marine Fisheries Scientific Collection Permit #153625.

## Data Archiving

Raw HiFi BAM and FASTQ for *Lycodicthys dearborni* and *Zoarces americanus* are available on NCBI SRA under the accession PRJNA1236397 [to be unembargoed upon acceptance]. HiFi FASTQ reads can be accessed on SRA for *Anarhichas lupus* (PRJNA980960) and *Anarhichas minor* (PRJNA982125). All code used for bioinformatic processing, analysis, and visualization is available on Github at https://github.com/om00sman/haplo_afp. *gfa_parser* is hosted and described at the Github repository https://github.com/snbogan/gfa_parser. *switch_error_screen* is hosted and described at the Github repository https://github.com/snbogan/switch_error_screen.

## Author Contributions

OWM led analysis and writing for the manuscript. SNB contributed to analysis and performed laboratory work. JLK and SNB contributed to writing. JLK and SNB conceived of the scope, aims, and analytical approach of the study. SNB wrote scripts for *gfa_parser* and *switch_error_screen* with consultation from OWM and JLK.

## Acknowledgements

This research was supported by National Science Foundation grant OPP-2312253 to JLK. We are grateful to Dr. Sean Place of Sonoma State University, CA, USA, for contributing samples and metadata from *Lycodichthys dearborni*. We thank Charles Jordan and Dr. James Gartland of the Virginia Institute of Marine Science, VA, USA, and the NorthEast Area Monitoring and Assessment Program for collecting samples of *Zoarces americanus*. We thank Dr. Muh-Ching Yee of the University of California, Santa Cruz, for her assistance with library preparation of *Z. americanus* DNA and comments on early drafts of the manuscript, Isabel Kline for her assistance in DNA extraction of *L. dearborni*, and the University of California, Davis, Genomics and Bioinformatics Core for performing HiFi sequencing and library preparation of *L. dearborni* DNA. We also thank Dr. Matthew Glasenapp and Kara Ryan for their comments on early drafts of the manuscript.

