## Supplemental Material for "Mitigating assembly and switch errors in phased genomes of polar fishes reveals haplotype diversity in copy number of antifreeze protein genes"

**Running title:** Haplotype diversity in antifreeze protein copy number

### 1. Supplemental Methods

*1.1. Tagmentation of Zoarces americanus DNA* - Barcoded primers were synthesized by IDT and resuspended at 100 µM in IDTE. Each barcoded primer contained a 5' universal PCR adapter sequence (5'-CGAACATGTAGCTGACTCAGGTCAC-3'), a 16 bp barcode, and a transposase mosaic end binding sequence (5'-AGATGTGTATAAGAGACAG-3'). Partially double-stranded adapters were formed by annealing 9 µL of 100 µM barcoded primer to 9 µL of a mosaic end reverse primer ([5'-PHO]CTGTCTCTTATACACATCT) with 2 µL of 0.5M NaCl in a 20 µL reaction. The mixture was incubated at 95°C for 5 minutes, cooled to 65°C at -0.1°C per second, incubated at 65°C for 5 minutes, and then cooled to 4°C at -0.1°C per second before storage at -20°C. Barcoded transposomes were assembled by incubating 2 µL of unloaded tagmentase (C01070010-20, Diagenode) with 2 µL of the annealed adapter at 23°C for 30 minutes. These transposomes were diluted in Tagmentase Dilution Buffer (C01070011, Diagenode) and titrated to working concentrations for optimal fragment sizes. Tagmentation reactions were performed on 1.5 µg of genomic DNA with 17 µL of 2X Tagmentation Buffer (C01019043-1000, Diagenode) and 2 µL of diluted barcoded transposome in a 34 µL final volume. The reaction was incubated at 55°C for 15 minutes, then stopped with 8.5 µL of 0.2% SDS and inactivated at 68°C for 5 minutes. Cleanup was performed using 21 µL (0.5X) of AxyPrep MAG PCR cleanup beads (Axygen MAGPCRCL500) to remove low molecular weight DNA and unincorporated adapters. The resulting DNA was used for PacBio SMRTbell library preparation.

*1.2. Bayesian modelling* - A Bayesian regression was used to test whether uncertainty in AFP copy number estimates increased in arrays of greater copy number. Percent uncertainty was modelled as a zero-inflated beta distribution using the R package brms v2.21.0 (Bürkner, 2017). The model was run with 40,000 MCMC chains and a warmup of 10,0000 chains. Significance in the beta parameter (slope) for the effect of median AFP copy number was assessed using a probability of direction test, a Bayesian corollary of the p-value that determines whether >95% of an effect's posterior distribution is greater than or less than 0 (Makowski *et al.*, 2019).

*1.3. Alternative modes for running* gfa_parser - We evaluated the compatibility of gfa_parser with GFA files produced by Verkko (Rautiainen *et al.*, 2023), minigraph (Li *et al.*, 2020), and Shasta (Shafin *et al.*, 2020) of by testing unphased and phased modes on GFAs from Verkko's reported assembly of the HG002 human genome (Rautiainen *et al.*, 2023), a minigraph pangenome graph of the GRCh38 reference genome previously used to evaluate gfabase, and a Shasta assembly of HG002 chromosome 21 reads previously used to evaluate gfabase (Lin). We evaluated errors, run time, and the successful formatting of FASTA outputs. *gfa_parser* can be run on GFA files output by Verkko, Shasta, minigraph, and hifiasm. The --format parameter can be set to the name of these four GFA assemblers to account for differences in formatting (see Supplemental Methods). Because Canu versions greater than 1.9 (Koren *et al.*, 2017) do not support GFA output, we did not test *gfa_parser* on GFAs from early versions of Canu.

*gfa_parser* can be run using two different classifications: unphased versus phased mode and standard versus Shasta and minigraph modes. Unphased mode takes in IDs for 5' and 3' start/end unitigs and automatically computes all directed acyclic paths and their contigs through the start and end. Phased mode requires a list of unitigs belonging to each haplotype, identifies all possible paths through the assembly of each haplotype, and outputs FASTA sequences for each path that are assigned to haplotypes. Shasta and minigraph modes are written to account for formatting differences in GFA files output by hifiasm and Verkko compared to Shasta and minigraph.

*1.4. Measuring genome-wide assembly uncertainty* - We sought to compare assembly uncertainty, defined as the total number of directed acyclic paths through a GFA, between AFP arrays and whole genomes. To achieve this, we developed a script (gw_paths.py) that expands on the approach of *gfa_parser* to compute the total number of paths at a genome-wide level. Dividing genome-wide paths by the number of unitigs in a GFA produces a normalized, whole-genome measure of assembly uncertainty that can be compared to uncertainty in specific regions. gw_paths.py first parses a GFA file into a directed graph using NetworkX (Hagberg *et al.*, 2008), where nodes represent unitigs and edges represent unitig overlaps. It then uses memoization, a caching technique that stores previously computed path counts to avoid redundant calculations (Cormen *et al.*, 2009), and depth-first traversal, a graph exploration method that follows each branch as far as possible before backtracking and computing the next longest path (Tarjan, 1974), to efficiently count all possible source-to-sink paths through a network of unitigs. 'Source-to-sink paths' begin at nodes with no incoming edges and end at nodes with no outgoing edges. These caching and traversing approaches support computational efficiency. gw_paths.py normalizes total paths by the number of unitigs in the GFA file. To handle extremely large integers, the total number of paths is calculated and reported on a log10 scale. A higher ratio of log10(paths)/unitig demonstrates increased uncertainty in the assembly of unitigs caused by unresolved *de novo* assembly and haplotype diversity.

After running gw_paths.py on all genomes in this study, we compared genome-wide assembly uncertainty to localized uncertainty values at AFP arrays. We predicted variation in assembly uncertainty as a function of whether measurements were made in AFP arrays versus whole genomes and a random effect of species identity using a generalized linear mixed model with a zero-inflated beta distribution fitted with glmmTMB v1.1.9 (Brooks *et al.*, 2017). We evaluated significance of the AFP vs. whole-genome effect using a likelihood ratio test comparing the test model to a null model lacking the AFP vs. whole-genome effect.

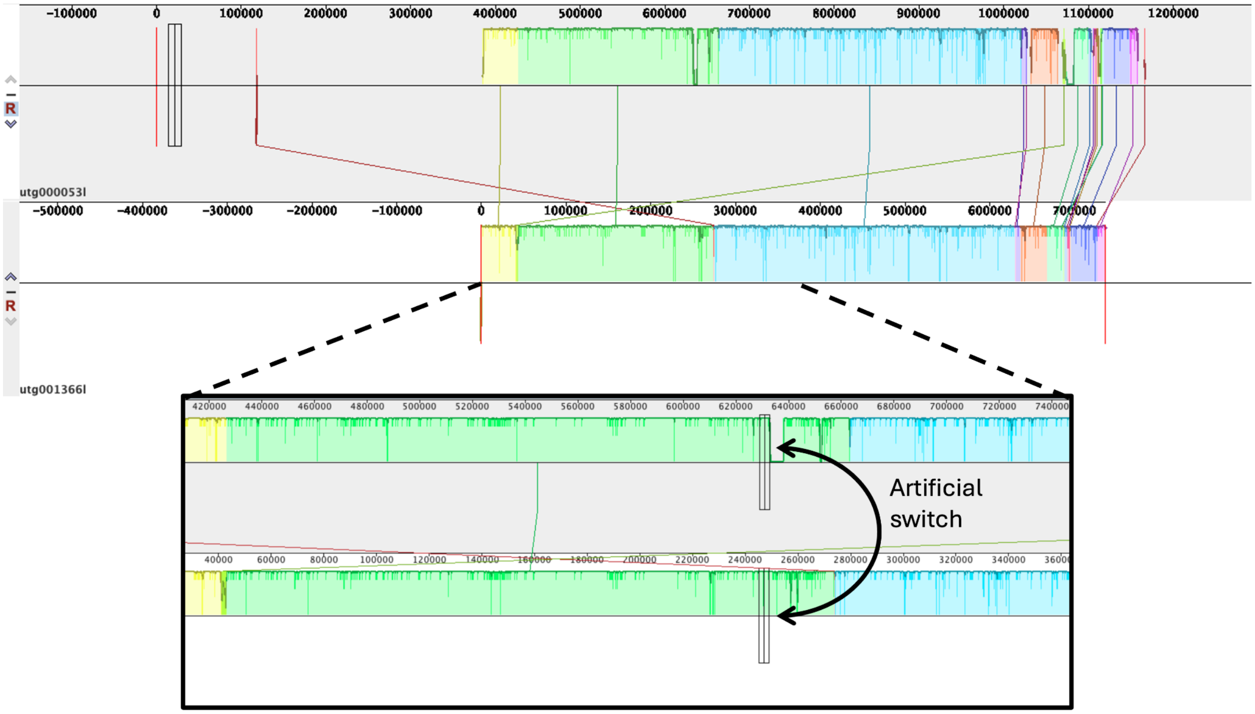


**Figure S1** | *Simulation of an artificial switch error in the Pholis gunnellus AFP array. A* MAUVE visualization of syntenic blocks conserved between *P. gunnellus* AFP array haplotypes is shown. Highlighted regions of the same color represent homologous sequences. Homologous blocks are also connected by lines. The height of each highlighted region represents sequence conservation level. The bottom of the figure zooms in on a region of high sequence divergence between homologous block 1 (yellow). A curve arrow depicts the position at which the 5' end of haplotype 1 (top track; gray background) was cut and concatenated with the 3' end of haplotype 2 (bottom track; white background). This artificial switch error was made to begin at a region of sequence divergence between haplotypes in order to impose polarized soft-clipping in HiFi reads that were realigned to the switch error region during realignment. In the presence of a switch error, it was expected polarized soft-clipping on one side of a switch error junction should arise during realignment. *switch_error_screen* was written to scan for this signature in BAM files and tested with the simulated switch error shown here.

### 2. Supplemental Results

*2.1. Genome-wide assembly uncertainty* - Analysis of GFA files by gw_paths.py demonstrated that assembly uncertainty was not significantly different between AFP arrays and whole genomes. gw_paths.py measured assembly uncertainty as the number of directed acyclic paths through unitigs of a GFA or GFA region, divided by the number of unitigs analyzed. The mean uncertainty in AFP arrays equaled 0.051 log10(paths) per unitig ± 0.029 SD. Mean genome-wide uncertainty equaled 0.130 ± 0.211 log10(paths) per unitig ($p = 0.8766$; Fig. S5).

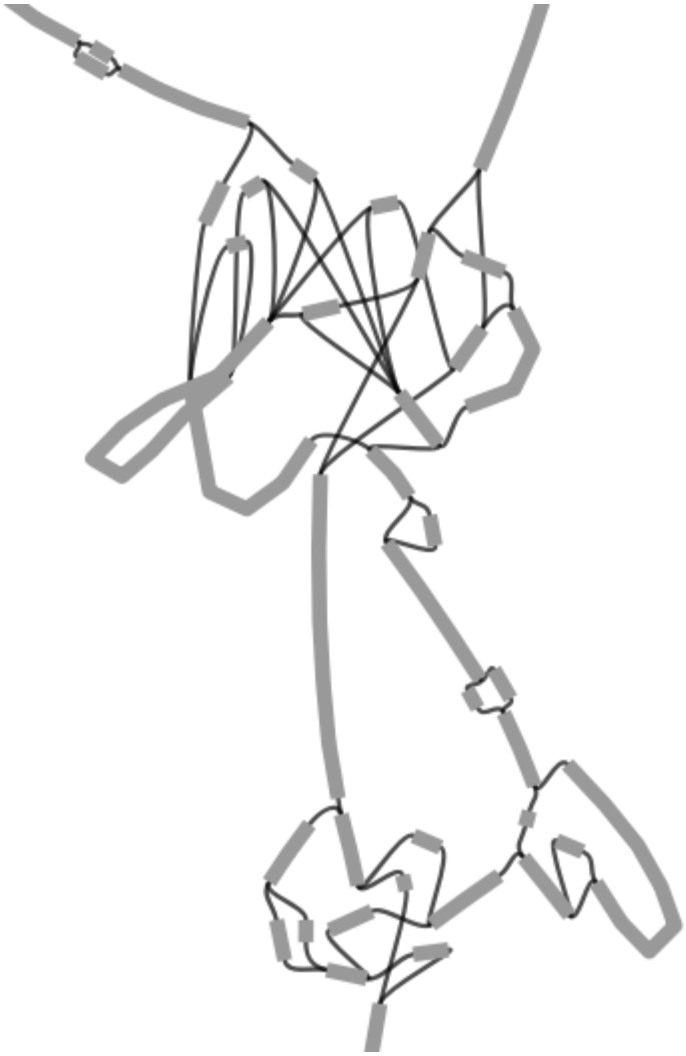

**Figure S2 |** *Unresolved haplotype structure in GFA file of* Leptoclinus maculatus *translocated AFP array 1.* An image exported from Bandage (Wick *et al.*, 2015) is shown that presents unitigs as thick gray lines (nodes) and overlaps between unitig sequences as thin black lines (edges). The range of all unitigs containing AFP genes in a translocated array is shown. The graphical fragment assembly (GFA) did not resolve to clear sets of paths that were non intersecting, which would have provided resolution of distinct haplotypes. For an example of resolved haplotypes in a GFA visualization, please refer to Figures 1 and 3 of the main text.

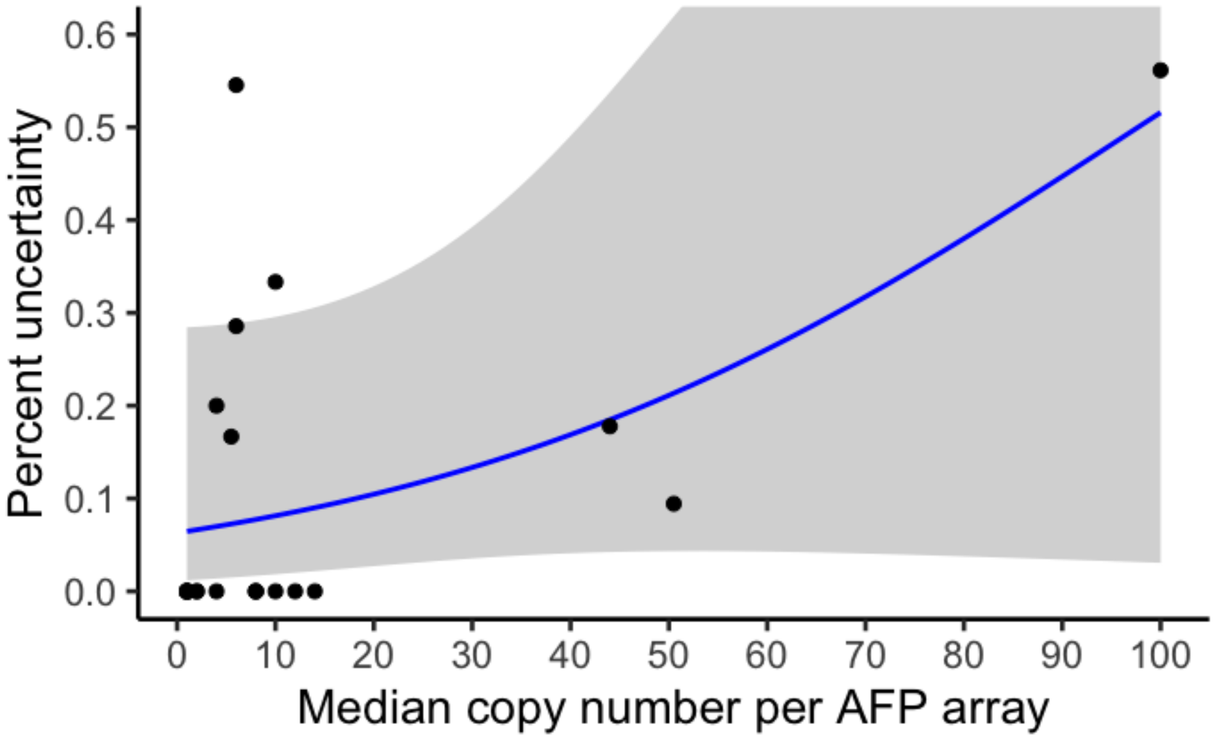


**Figure S3 |** *Percent uncertainty across increasing AFP copy number.* Median copy number of each AFP array in all species plotted against percent uncertainty for that array. Percent uncertainty was calculated as the difference in copy number between the assemblies with the minimum and maximum AFP copy numbers divided by the maximum copy number.

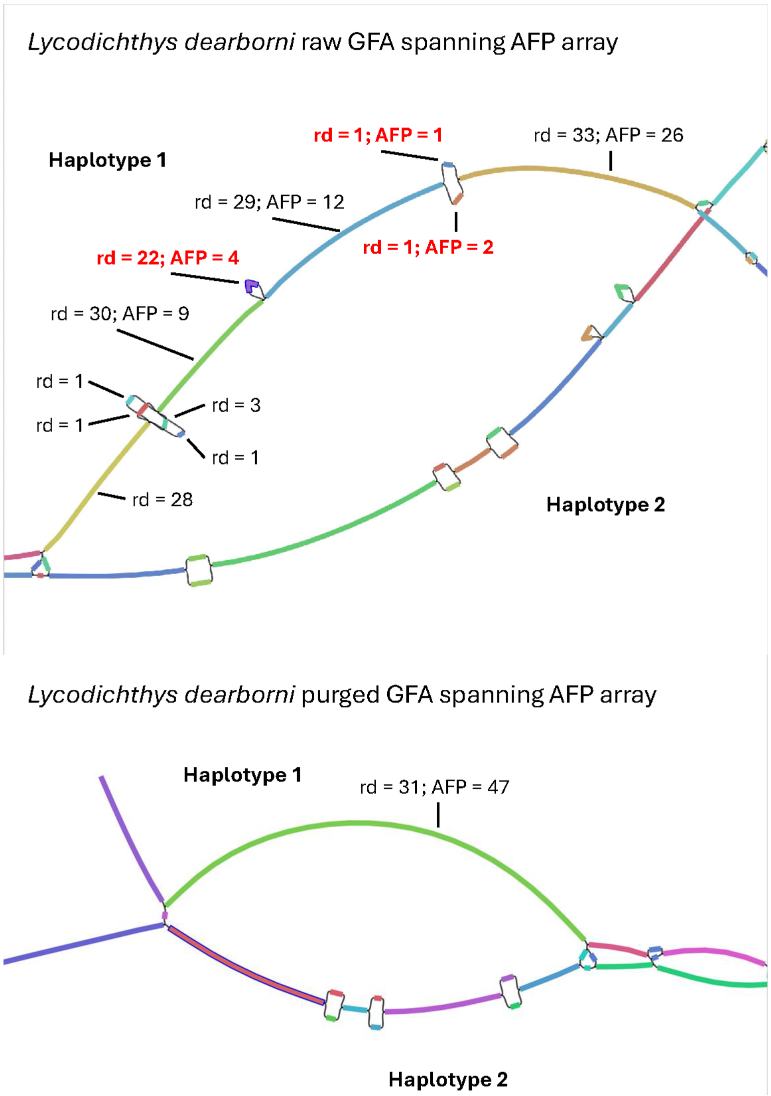


**Figure S4** | *An example of AFP gene loss during small bubble popping.* Two hifiasm GFA files are visualized as rendered in Bandage. (Top) A raw GFA from *Lycodicthys dearborni* spanning the AFP array is shown. Unitigs are represented as thick colored lines. Overlaps in unitig sequence are represented by thin black edges. The mean read depth (rd) of each haplotype 1 unitig is annotated. If haplotype 1 unitigs contained intact AFP genes, AFP copy number was annotated alongside read depth. Unitigs of low-to-moderate read depth that contained AFP genes and were removed from the GFA during bubble popping are associated with bold, red annotations. (Bottom) A purged GFA file is shown that underwent bubble popping and duplicate purging. Haplotype 1 unitig read depth and AFP copy number are annotated as in the raw GFA.

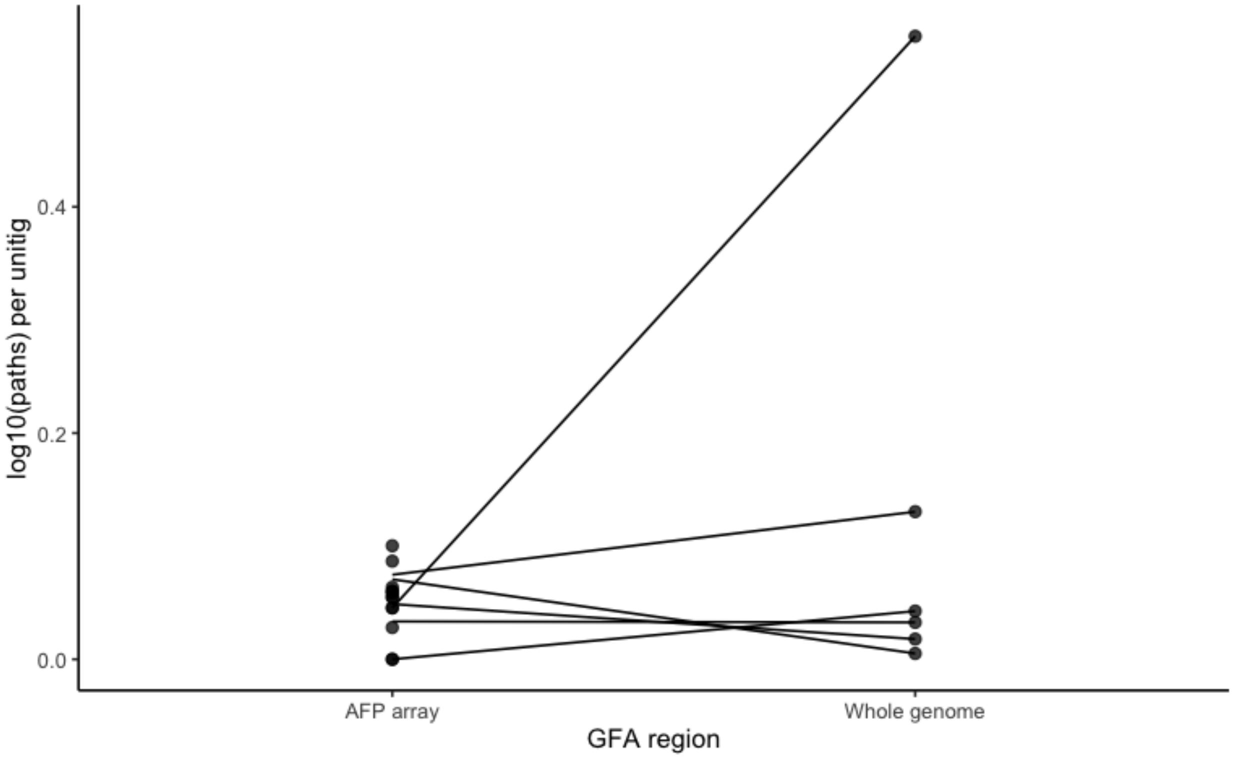


**Figure S5** | *Assembly uncertainty in AFP arrays versus whole genomes*. The y-axis represents the log10 of the number of directed acrylic paths through GFA files divided by the number of unitigs in those paths. log10(paths) per unitig was measured in subsetted GFAs at AFP array loci and in complete GFA files at a whole-genome scale. Along the x-axis, points above "AFP array" represent uncertainty at individual AFP arrays. Lines connect the mean uncertainty at all AFP arrays in a species to that species' whole-genome uncertainty measure.
